# Low-pass nanopore sequencing for measurement of global methylation levels in plants

**DOI:** 10.1101/2024.08.12.607255

**Authors:** Yusmiati Liau, Annabel Whibley, Amy Hill, Bhanupratap Vanga, Meeghan Pither-Joyce, Elena Hilario, Sarah Bailey, Susan J. Thomson, Darrell Lizamore

## Abstract

Nanopore sequencing enables detection of DNA methylation at the same time as identification of canonical sequence. A recent study validated low pass nanopore sequencing to accurately estimate global methylation levels in vertebrates with sequencing coverage as low as 0.01x. We investigated the applicability of this approach to plants by testing three plant species and analysed the effect of technical and biological parameters on estimate precision and accuracy. Our results indicate that a higher coverage (0.1x) is required to assess plant global methylation at an equivalent accuracy to vertebrates. Shorter read length and a closer sequence match between sample and reference genome improved measurement accuracy. Application of this method in *Vitis vinifera* showed consistent global methylation levels across different leaf sizes, and different sample preservation and DNA extraction methods, whereas different varieties and tissue types did exhibit methylation differences. Similarly, distinct methylation patterns could be observed in different genomic features. Our findings suggest the suitability of this method as a low-cost screening tool for validation of experimental parameters, developmental time courses and to assess methylation status for different modification types and sequence contexts at the level of whole genome or for abundant genomic features such as transposable elements.

## Introduction

Skimseq, or genome skimming, is defined as untargeted, low-pass sequencing, usually at lower than 2x coverage (Hu et al., 2023). Whilst originally employed as an approach to comprehensively capture over-represented elements within the sample, such as organelle, viral or parasitic genomes (Ripma et al., 2014; Weitemier et al., 2014), this method can also provide reliable, cost-effective estimates of global genomic parameters (e.g. for investigation of highly abundant transposable elements (Lwin et al., 2017)), or it can be combined with genomic imputation for high-throughput genotyping by sequencing (Kumar et al., 2021).

The Oxford Nanopore Technologies sequencing platform reports not only canonical bases but also native DNA modifications including methylated and hydroxymethylated cytosines (5mC and 5hmC, respectively) and methylated adenosines (6mA) and, consequently, genomic insights can extend from the canonical sequence to the epigenetic properties of the samples (Laszlo et al., 2013; Schreiber et al., 2013; Simpson et al., 2017). Using low coverage nanopore sequencing for methylation detection in vertebrate genomes, Faulk reported the high precision and accuracy of global methylation assessment at only 0.01x coverage (i.e. 30 Mb per sample) (Faulk, 2023). The report also demonstrated the accuracy of methylation level estimation for *Alu* transposon elements at 0.001x (i.e. 3Mb per sample). The approach was shown to be reproducible across technical and biological replicates and was reportedly not affected either by read length or quality.

In vertebrates, cytosines adjacent to guanine (CG) can be methylated by DNA Methyltransferases, either during DNA replication or during early development (Klughammer et al., 2023). In contrast, cytosines in plant genomes can be methylated in a variety of sequence contexts, which are mediated by different enzymatic pathways. These methylation contexts are categorised as CG, CHG or CHH, where H is either A, T, or C. Methylation level is typically highest in CG, followed by CHG and CHH contexts, with wide variation throughout different plant species (Niederhuth et al., 2016). Methylation in the CG context is the main methylation found in gene bodies and is regulated by METHYLTRANSFERASE 1 (MET1), while CHG methylation is largely associated with repetitive sequences, with the methylation in this motif being copied to the newly synthesised strand by CHROMOMETHYLASE 3 (CMT3) during DNA replication. In contrast, CHH methylation is not symmetrical, and therefore must be applied in a sequence-guided manner. This is achieved by CMT2, which targets heterochromatic DNA, or by DOMAINS REARRANGED METHYLTRANSFERASE 2 (DRM2) during RNA-dependent DNA methylation (Liu et al., 2023; Zhang et al., 2018a). Adenosine methylation (6mA) has also been observed in low levels in plants, enriched in genic regions and, in contract to 5mC, has been shown to be positively associated with gene expression (Zhang et al., 2018b; Zhang et al., 2023). Exploring genome methylation by context therefore gives important biological insight into chromatin structure and transposon silencing, in addition to gene regulation.

To investigate how effectively skimseq can be applied to study global methylation in plants, we tested the precision and accuracy of the approach in nanopore sequencing datasets from three plant species: *Vitis vinifera* (grapevine), *Arabidopsis thaliana* and *Actinidia melanandra* (purple kiwifruit). We examined the influence of technical and biological factors such as read length, methylation entropy, genetic heterozygosity, genome size, and reference genome choice on the accuracy of this approach for measuring global methylation.

Having established appropriate coverage thresholds for skimseq in plants, we used this approach to investigate the variation of global methylation levels with respect to sample preservation and DNA extraction methods, as well as across different grapevine tissues and varieties. Sample preservation method is an important factor for plant genomic analysis, to ensure good quality of nucleic acid and preservation of biological information. For genomic analysis, suitable DNA can normally be obtained from samples collected without immediate freezing. In contrast, samples are typically snap-frozen in the field using liquid nitrogen or immersed in RNA-preserving chemicals such as RNALater to ensure high-quality RNA can be extracted for transcriptomic studies. Scant data is available regarding the effect of sample preservation methods on the stability of DNA methylation. To address this, we used skimseq to compare global methylation levels of tissue samples collected using four different methods (snap-frozen in liquid nitrogen, frozen with dry ice, packaged with silica-gel, and stored at room temperature), as well as two different DNA extraction methods. To compare the impact of technical methods with true biological variation, we also compared the global methylation level between different grapevine tissues of the same variety, and different grapevine varieties.

## Methods

### Samples

*Vitis vinifera* cv. ‘Sauvignon Blanc’, clone UCD1 (FPMS1) young leaf samples were collected from the New Zealand Winegrowers National Vine Collection held at Lincoln University. Vine-harvested leaves of *Actinidia melanandra* (ME02_01) were collected from Te Puke, Bay of Plenty. Leaf punches of *Arabidopsis thaliana Col-0* ecotype were obtained from a single lab-grown plant. All samples were snap-frozen by immersion in liquid nitrogen at the time of collection.

To compare global methylation levels between technical methods, tissue types and cultivars, leaf and tendril samples were collected from *Vitis vinifera* cv. Sauvignon Blanc, clone UCD1 (FPMS1) grafted onto rootstock 3309 from a commercial vineyard (Waiata Vineyard, Tiki Wine) in North Canterbury, New Zealand. Samples were snap-frozen in the field and stored at -80 °C until DNA extraction, except for some samples that were specifically collected for assessment of the impact of preservation methods on sequence properties. These alternative preservation methods included: (i) collection into silica-gel, whereby leaves were collected into an empty teabag and put inside a Ziploc bag with 30g of silica gel, refrigerated at 2-8 °C overnight before being stored at -80 °C; (ii) frozen by packaging in dry ice; (iii) a room temperature condition, where leaves were collected without any cooling method, and left at room temperature for 2 hours before storage at -80 °C. Two other *Vitis vinifera* varieties, Pinot Gris and Pinot Noir were also collected using the liquid nitrogen sampling method. Two to three replicates were collected for each set of experimental parameters.

### DNA extraction

Frozen tissues were ground in liquid nitrogen using a mortar and pestle or homogenised in 2 mL tubes using a TissueLyzer instrument (Qiagen) immediately prior to DNA extraction. Purified DNA for the *Vitis vinifera* high-coverage sample and *Actinidia melanandra* were extracted using the Nucleobond High Molecular Weight DNA kit (Macherey-Nagel, Düren, Germany). Nuclei isolation was performed prior to DNA extraction for the *Actinidia* sample using the PacBio protocol (Pacific Biosciences, 2022). Size selection to remove short reads was performed for both DNA extracts using the Short Read Eliminator XL reagent (Pacific Biosciences, CA, USA). The *Arabidopsis* genomic DNA was extracted using a CTAB - based protocol in which a leaf punch was incubated for 2 hours at 56 °C in CTAB buffer (as described in (Hilario, 2018) with gentle homogenisation of the tissue during incubation, followed by one round of chloroform:isoamyl alcohol (24:1) purification, ethanol precipitation of the nucleic acids and resuspension in 1X TE buffer (pH 7.5). RNase treatment was performed after extraction.

For samples intended for comparison of different *Vitis vinifera* tissue types, varieties and pre-analytical methods, DNA was extracted using the Nucleomag plant DNA kit (Macherey-Nagel, Düren, Germany), a CTAB-based extraction protocol, with the purification step automated on Eppendorf EpMotion 5075 liquid-handling robot. Three samples were also re-extracted using an alternative SDS-based extraction method (Russo, 2020). DNA concentrations were measured using the Qubit broad range kit on a Qubit Flex instrument and purity was determined using a nanodrop 8000 (both from Thermo Fisher Scientific, Waltham, MA, USA).

### Library preparation

Sequencing libraries for the grapevine, *Arabidopsis* and kiwifruit samples were prepared using the ligation sequencing kit from Oxford Nanopore Technologies (SQK-LSK114) following the manufacturer’s protocol and sequenced on separate R10.4.1 flow cells. For the extended *Vitis vinifera* population samples, barcoding and sequencing library preparation was performed using the Oxford Nanopore Rapid barcoding kit V14 (SQK-RBK114.96) following the manufacturer’s protocol and sequenced across two R10.4.1 flow cells. All sequencing was performed on an PromethION P24 instrument (Oxford Nanopore Technologies) at the Bragato Research Institute (Lincoln).

### Data analysis

The publicly available dataset from the Oxford Nanopore Open Data Project human cell line GM12878 (HG001; Genome in A Bottle Consortium) was used as a comparison to the plant datasets generated in this study (see https://labs.epi2me.io/giab-2023.05/). Raw Fast5 sequence data files were converted into pod5 where necessary using pod5 tool v0.2.4 (https://github.com/nanoporetech/pod5-file-format), and re-basecalled using dorado v0.3.2 (https://github.com/nanoporetech/dorado) with the ‘super accurate’ (SUP) basecalling model and modified base models for both 5mC and 6mA (dna_r10.4.1_e8.2_400bps_sup@v4.2.0_5mC@v2 and dna_r10.4.1_e8.2_400bps_sup@v4.2.0_6mA@v2) in all contexts. The resulting BAM files were converted back to fastq, with modification tags preserved, using SAMtools v1.0 (RRID:SCR_002105) (Danecek et al., 2021). Quality filtering of the reads (minimum Phred average quality score of 10) was performed using chopper v0.5.0 (https://github.com/wdecoster/chopper) and these reads, containing MM/ML tags were mapped to reference genomes using minimap2 v2.26 (RRID:SCR_018550) (Li, 2021) with the *-ax map-ont* presets. Reference genomes used were PN40024.v4 and SB1031v1 for *Vitis vinifera*, TAIR10 (GenBank accession: GCA_000001735.2) for *Arabidopsis thaliana*, ME02_01 v2.5 for *Actinidia* melanandra, and GRCh38 (GenBank accession: GCA_000001405.15) for human dataset.

Coverage was assessed using Mosdepth v0.3.3 (RRID:SCR_018929) (https://github.com/brentp/mosdepth) and was used to calculate the proportion of a dataset required to downsample the mapped BAM files to the desired coverage levels using SAMtools v1.0 (RRID:SCR_002105) (Danecek et al., 2021). Modkit v0.1.8 (https://github.com/nanoporetech/modkit) was used to process the BAM files to generate, filter and process methylation calls, producing a BEDmethyl output file, and to generate reference BED files containing genomic position of CG, CHG, CHH, and 6mA contexts.

Grapevine gene annotations were downloaded from https://integrape.eu/ and transposable element (TE) regions were annotated using EDTA v2.1.0 (RRID:SCR_022063) (Ou et al., 2019). No curated library was provided to EDTA, *de novo* element discovery with RepeatModeler2 was enabled, and the coding sequences from the PN40024.v4 assembly release were provided to limit misclassification of genes as transposable elements. Methylation data on each region and context were generated using BEDtools v2.29.2 (RRID:SCR_006646) (Quinlan and Hall, 2010). For analysis of data with different read lengths, reads were grouped based on read length criteria using Chopper. As the Vitis library contains mainly long reads, the 5kb length data was generated by trimming the reads to this length using reformat.sh from bbmap v39.01 (RRID:SCR_016965) (Bushnell, 2014). Phasing and separation of reads to each haplotype was performed using WhatsHap v1.6 ((Martin et al., 2016). Global methylation levels were calculated using AWK scripts, as described by (Faulk, 2023). Error rate was calculated as a mean difference in percentage between the highest and lowest value of the 10 replicates compared to true value, i.e. value obtained from data with high coverage. Nanoplot v1.41.0 (RRID:SCR_024128) was used to generate sequencing metrics such as read length and quality (De Coster and Rademakers, 2023). Methylation entropy was calculated using DMEAS (He et al., 2013) on data with ∼10x coverage. Differences of methylation levels among groups were analysed using one-way ANOVA (RRID:SCR_002427) followed by a Tukey’s multiple comparison test and Plots were created in using ggplot2 (RRID:SCR_014601) (Wickham, 2009) in R v4.2.2. All bioinformatics analysis was performed with the aid of New Zealand eScience Infrastructure (NeSI) high performance computing facilities.

## Results

### Performance of skimseq approach for global methylation assessment in grapevine

We sequenced one grapevine sample to a total depth of 168x, and downsampled this dataset to 10x, 1x, 0.1x, 0.01x and 0.001x coverage, with ten bootstrap replicates performed at each coverage level. Using the analysis approach described in (Faulk, 2023), global methylation level estimates for CG, CHG, CHH and 6mA contexts were computed for each coverage level. In all sequence contexts, global estimates of methylation level were consistent with the original value down to coverage of 0.1x, with error rate <5% for CG and 6mA. Error rates were slightly higher for CHG and CHH contexts, (<10% at 0.1x and <5% at 1x). These observed error rates were higher than those previously reported in vertebrates (Faulks, 2023), especially for non-CG contexts (Table 1 and Figure 1).

**Table 1.** Summary statistics for global methylation level estimates in CG, CHG, CHH, and 6mA contexts in the full dataset and at downsampled coverage from 10x to 0.001x for *Vitis vinifera*, with 10 bootstraps performed for each subsample.

| Coverage | 168x | 10x | 1x | 0.1x | 0.01x | 0.001x |
| --- | --- | --- | --- | --- | --- | --- |
| **CG** |  |  |  |  |  |  |
| Context |  |  |  |  |  |  |
| Mean | 61.53 | 61.61 | 61.72 | 61.17 | 62.64 | 59.67 |
| Max |  | 61.72 | 62.47 | 62.22 | 65.54 | 76.97 |
| Min |  | 61.45 | 61.06 | 59.76 | 57.77 | 44.51 |
| SD |  | 0.1 | 0.38 | 0.84 | 2.91 | 9.36 |
| SE |  | 0.03 | 0.12 | 0.26 | 0.92 | 2.96 |
| Upper error % |  | 0.3 | 1.52 | 1.12 | 6.52 | 25.09 |
| Lower error % |  | -0.14 | -0.77 | -2.88 | -6.12 | -27.67 |
| Mean error % |  | 0.22 | 1.15 | 2.00 | 6.32 | 26.38 |
| **CHG** |  |  |  |  |  |  |
| Context |  |  |  |  |  |  |
| Mean | 27.24 | 27.32 | 27.44 | 26.97 | 28.05 | 27.64 |
| Max |  | 27.45 | 28.03 | 28.7 | 31.67 | 43.88 |
| Min |  | 27.1 | 26.86 | 24.91 | 22.55 | 18.3 |
| SD |  | 0.11 | 0.35 | 1.31 | 3.07 | 7.94 |
| SE |  | 0.03 | 0.11 | 0.42 | 0.97 | 2.51 |
| Upper error % |  | 0.77 | 2.9 | 5.38 | 16.26 | 61.09 |
| Lower error % |  | -0.51 | -1.39 | -8.56 | -17.22 | -32.82 |
| Mean error % |  | 0.64 | 2.14 | 6.97 | 16.74 | 46.96 |
| **CHH** |  |  |  |  |  |  |
| Context |  |  |  |  |  |  |
| Mean | 1.84 | 1.84 | 1.85 | 1.82 | 1.93 | 1.66 |
| Max |  | 1.85 | 1.88 | 1.91 | 2.28 | 2.04 |
| Min |  | 1.84 | 1.81 | 1.67 | 1.54 | 1.14 |
| SD |  | 0 | 0.02 | 0.08 | 0.22 | 0.27 |
| SE |  | 0 | 0.01 | 0.02 | 0.07 | 0.08 |
| Upper error % |  | 0.76 | 2.52 | 4.07 | 24.21 | 11.09 |
| Lower error % |  | -0.04 | -1.32 | -9.06 | -16.09 | -37.73 |
| Mean error % |  | 0.40 | 1.15 | 2.00 | 6.32 | 26.38 |
| **6mA** |  |  |  |  |  |  |
| Context |  |  |  |  |  |  |
| Mean | 4.1 | 4.11 | 4.12 | 4.08 | 4.2 | 3.99 |
| Max |  | 4.12 | 4.17 | 4.2 | 4.59 | 4.82 |
| Min |  | 4.09 | 4.08 | 3.91 | 3.58 | 3.25 |
| SD |  | 0.01 | 0.03 | 0.11 | 0.33 | 0.56 |
| SE |  | 0 | 0.01 | 0.03 | 0.11 | 0.18 |
| Upper error % |  | 0.47 | 1.72 | 2.45 | 12.02 | 17.62 |
| Lower error % |  | -0.18 | -0.62 | -4.59 | -12.7 | -20.67 |
| Mean error % |  | 0.32 | 1.17 | 3.52 | 12.36 | 19.14 |

**Figure 1.**
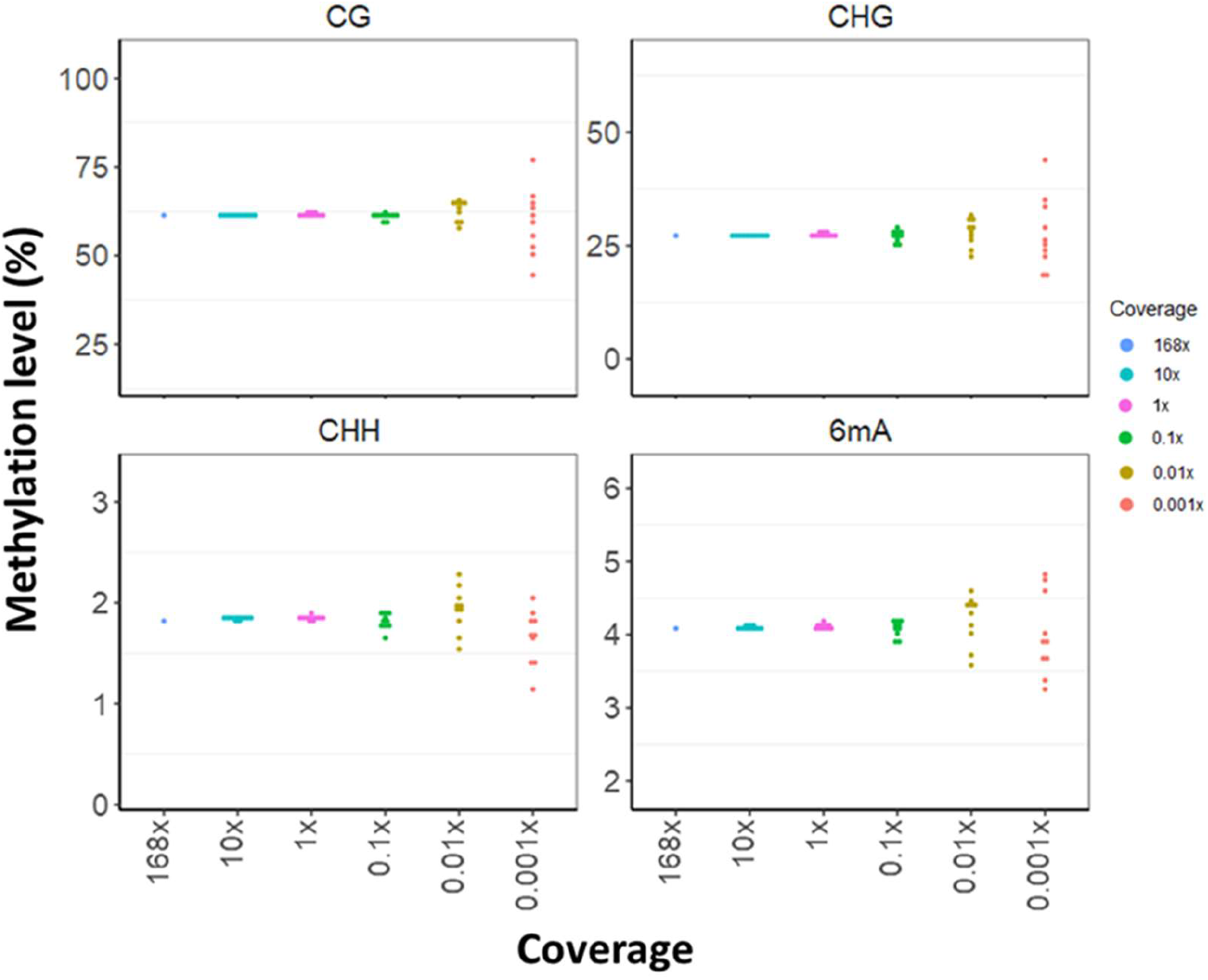
Global methylation level estimates in CG, CHG, CHH, and 6mA contexts in the full dataset and at downsampled coverage from 10x to 0.001x for *Vitis vinifera*, with 10 bootstraps performed for each subsample.

### Comparison of different plant data

To determine whether the higher error rates of skimseq global methylation levels in grapevine, relative to that reported in vertebrates, is common to other plant species, we extended our analyses to two other species: *Arabidopsis thaliana*, sequenced to ∼12x, and *Actinidia melanandra*, sequenced to ∼70x coverage. We also analysed a control human dataset, sequenced to 11x coverage using the same flow cell type and library kit chemistry as our plant datasets. These three additional samples were downsampled using the same approach, and error rates of global methylation in CG context were compared. Data from the human sample showed a similar pattern to that reported by Faulk (2023), with accurate estimation of methylation level down to 0.01x (error approximately 3%). Despite a large difference in absolute levels of CG methylation (30% compared to 65%), *Arabidopsis thaliana* showed a similar error profile to *Vitis vinifera* across all coverage levels, with an error rate of >5% at 0.1x. *Actinidia melanandra* showed a lower error rate compared to *Vitis vinifera* and *Arabidopsis thaliana* at 0.1x and 0.01x (0.31% and 3% respectively), but a similar error rate at 0.001x. The methylation entropy also differs notably among the three plant species (Figure 2 and Table S1).

**Figure 2.**
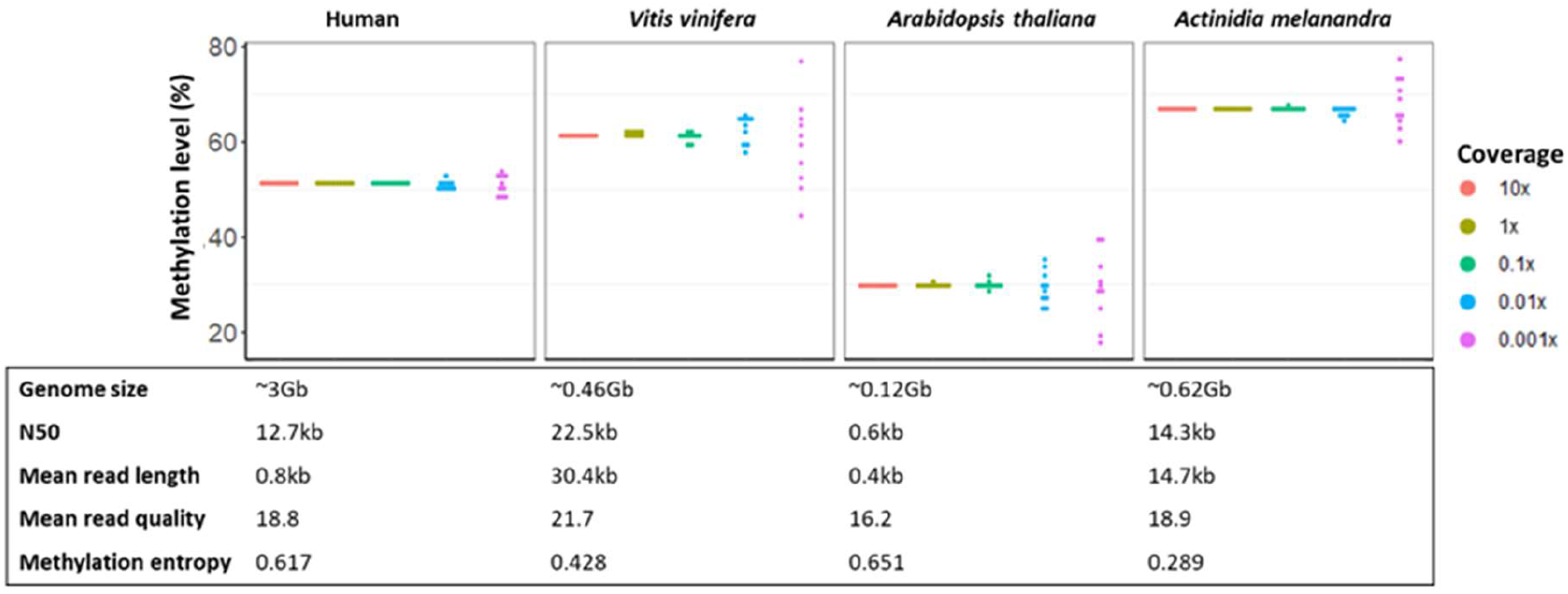
Global methylation level estimates of human, *Vitis vinifera, Arabidopsis thaliana*, and *Actinidia melanandra* datasets in CG context at coverage from 10x to 0.001x, with 10 bootstraps performed for each subsample.

The four datasets vary in terms of library properties (e.g. read length distributions), reference genome properties (e.g. reference assembly quality and the degree of matching between the sample and the reference assembly) and in genome biology (e.g. levels of heterozygosity), although all are from diploid species. We sought to understand how these features might contribute to the variation of error rate (Figure 3 and Table S2-4).

**Figure 3.**
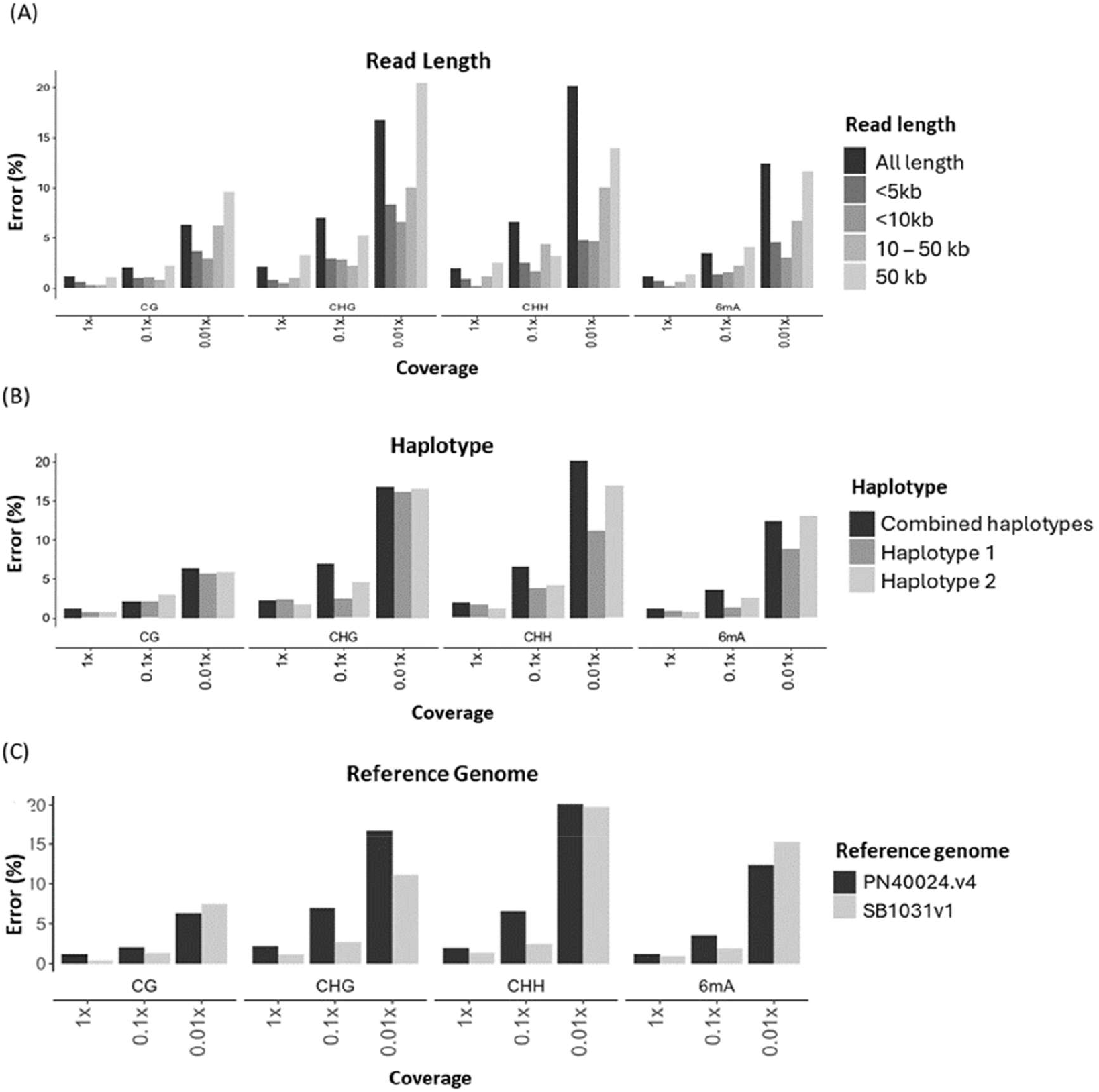
Variation of error rates based on biological and technical factors: (A) Error rates of Vitis datasets with different read length, (B) error rates of Vitis datasets in separate haplotypes versus combined haplotypes, (C) error rates of Vitis datasets mapped to two different reference genomes.

To compare the effect of library read length on accuracy, we grouped the *Vitis* dataset into four different length ranges (5kb, <10kb, 10 to 50 kb, and >50 kb), and performed downsampling separately on each group. The error rates were considerably lower in datasets with shorter reads, especially for CHG and CHH contexts (Figure 3A).

Plant genomes can be highly heterozygous (Claros et al., 2012), for example *Vitis vinifera* genomes have up to 13% sequence divergence between haplotypes (Jaillon et al., 2007). To account for the effect of this genetic heterogeneity, we partitioned the reads by haplotype and downsampled alignments each containing a single haplotype separately. No difference was observed between error rates of each haplotype and that of the combined data (Figure 3B).

Lastly, we observed lower error rates in kiwifruit, which was mapped to an in-house reference genome built using reads from the exact same sample, while the grapevine sample was initially mapped to the commonly used Vitis reference genome, PN40024.v4, which was built from a different *Vitis vinifera* variety (Velt et al., 2023). We re-mapped the grapevine sample onto our in-house reference genome, built using data from this exact Vitis sample, and re-performed the downsampling. This resulted in lower error rates, especially for the CHG and CHH contexts at 0.1x coverage (Figure 3C).

### Performance of skimseq grapevine methylation level in different genomic features

Methylation levels vary in different genomic contexts and regions (Figure 4A). CG methylation levels are relatively high throughout the genome, while CHG methylation levels are relatively high across transposable elements (TE) but lower in genic regions. In the *Vitis* sample, TE and genic regions comprised ∼45% and ∼33% of the genome, respectively. To determine the precision and accuracy of skimseq approach for estimating methylation levels in these genomic regions across different sequence contexts, we annotated the BEDmethyl files with region information and calculated the methylation levels in respective regions. The error rates of methylation levels in TE and genic regions at 0.1x to 10x coverage are comparable or lower than genome-wide assessment, except for CHG methylation in genic regions and CHH and 6mA methylation at separate classes of TE regions (Figure 4, Figure S1, and Table S5).

**Figure 4.**
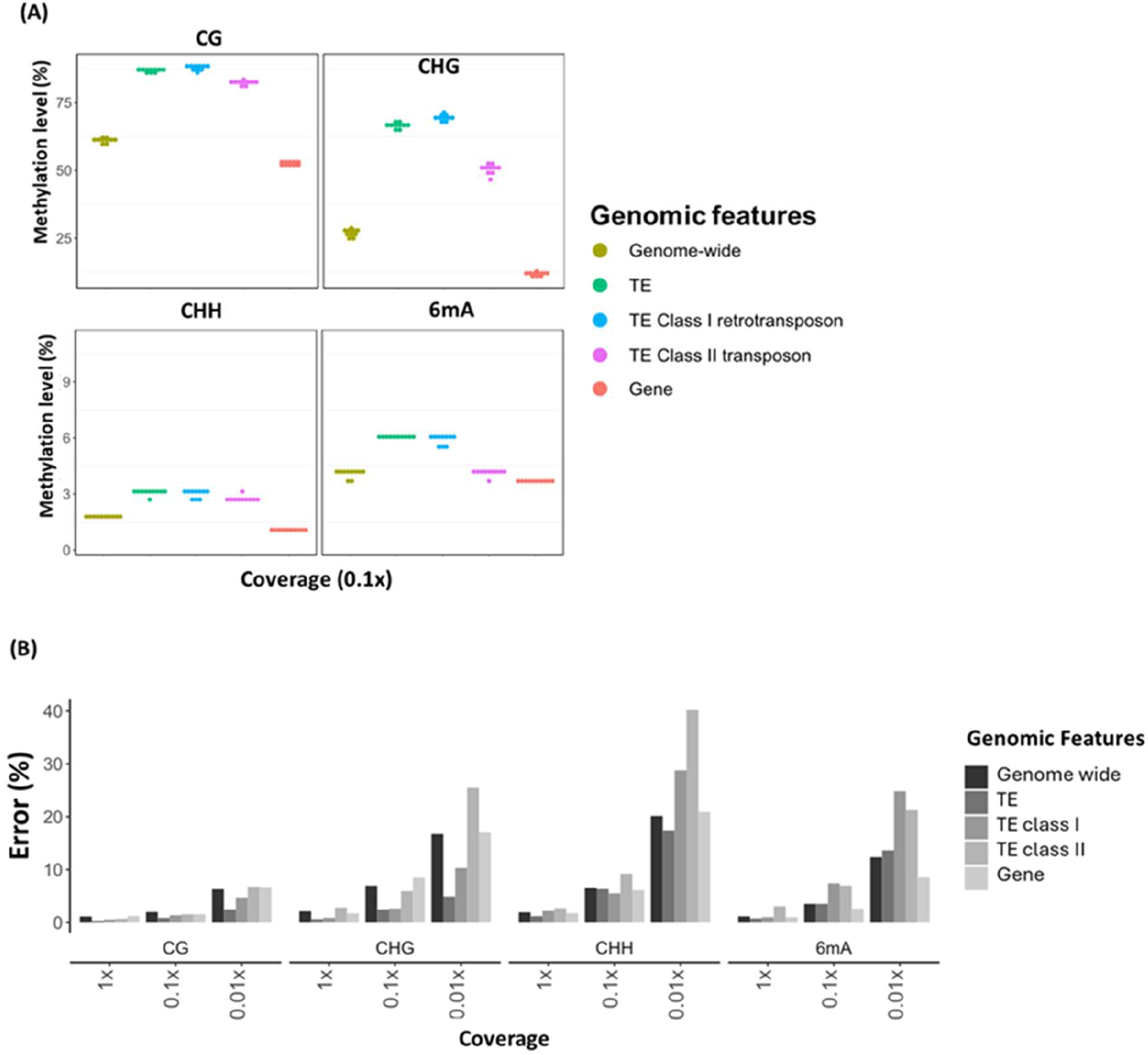
*Vitis vinifera* methylation level and error rates in different genomic features. (A) Methylation levels in different genomic contexts and features at coverage of 0.1x, (B) Error rates at coverage of 1x to 0.01x.

### Skimseq for methylation measurement of different grapevine tissue preservation methods

Having established the coverage threshold (0.1x) necessary to measure global methylation, as well as for TE or genic regions, we applied this approach to validate the consistency of DNA methylation levels at low coverage across different tissue types, and among different preservation and extraction methods in *Vitis vinifera* cv. Sauvignon Blanc. In addition, we also assessed the methylation level in a second grapevine variety, cv. ‘Pinot Noir’ and its clonal variant, ‘Pinot Gris’. Samples were sequenced to 0.24x – 2.31x coverage with N50 ranging from 2.9 - 9.9kb (Table S6).

Analysis of these data showed methylation measurements to be independent of these parameters, with some exceptions. Global CG methylation levels did not significantly differ across tissue types, preservation methods, or DNA extraction methods. Global methylation of ‘Sauvignon Blanc’ appears lower than the ‘Pinot’ varieties but did not cross the threshold for significance (Figure 5).

**Figure 5.**
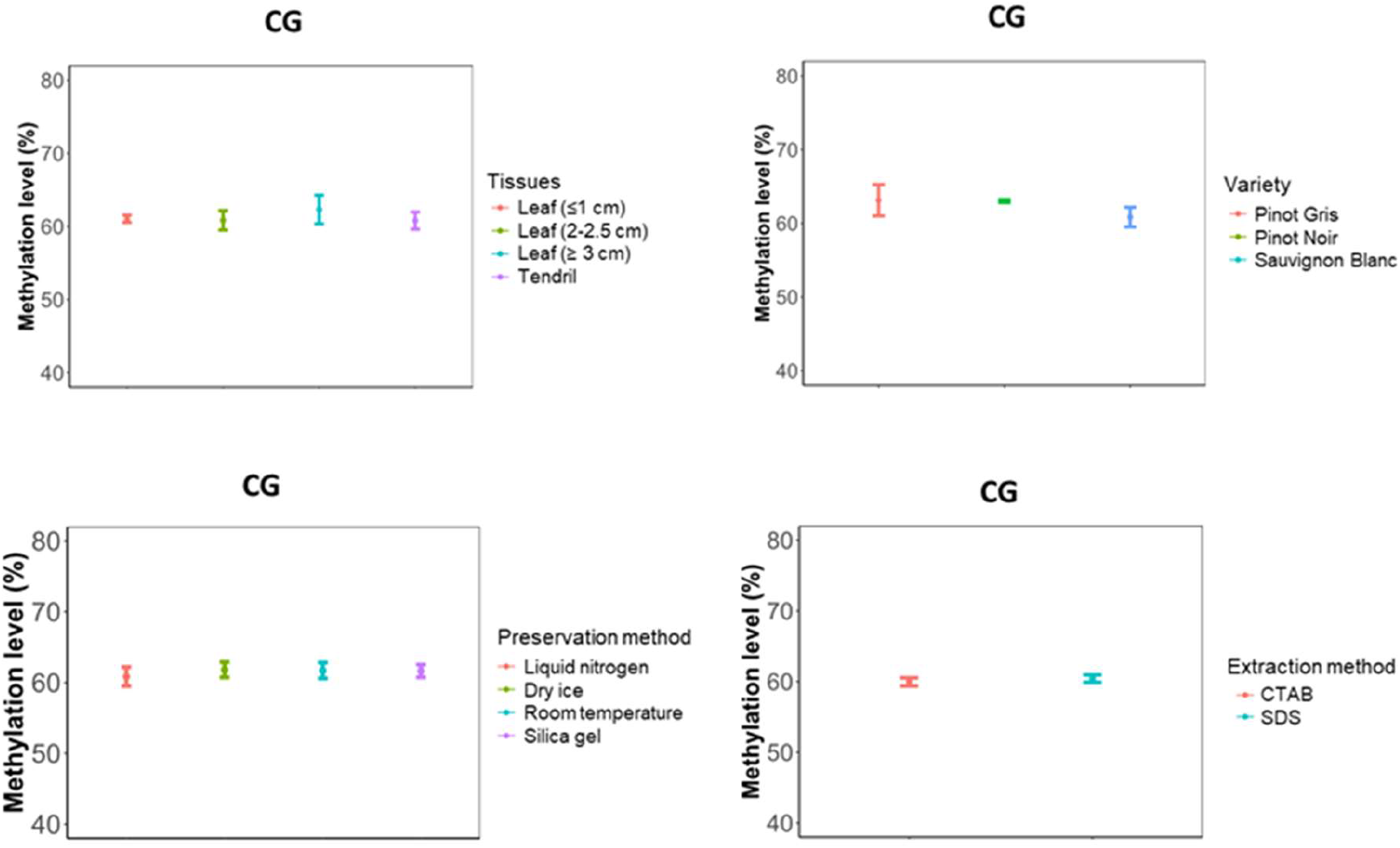
*Vitis vinifera* global methylation levels (mean ± sd) in CG context across different tissues, varieties, sample preservation methods, and DNA extraction method. CTAB = Cetyltrimethylammonium bromide, SDS = Sodium dodecyl sulfate.

We also showed that methylation levels were consistent between three different sizes of young leaves (≤1 cm, 2-2.5cm, and ≥3 cm) and tendril tissue for CG, CHG and 6mA contexts, while methylation in CHH context were higher in tendril compared with the two smaller leaf sizes (Table S6 and Figure S2).

Interestingly, the two different *Vitis* varieties included in this study did differ in CG methylation with regards to class II TEs. This feature type showed significantly higher methylation levels in Pinot varieties compared with Sauvignon Blanc (Pinot Gris=83.17±0.59%, Pinot Noir=82.19±0.29%, Sauvignon Blanc=79.75±0.97%) (Figure S3).

Lastly, we showed that methylation levels were independent of sample preservation methods (liquid nitrogen, dry ice, silica-gel and room temperature) and extraction methods (CTAB-based versus SDS-based) (Figure S4 and S5).

## Discussion

Various methods have been developed to measure DNA methylation. Genome-wide or targeted methylation at a single-base level can be measured using bisulfite treatment followed by whole genome or targeted sequencing. At a lower resolution, methods such as methylation-sensitive amplification polymorphism offer more affordable ways to investigate DNA methylation status for specific sequence motifs (Agius et al., 2023). Global methylation status can be assessed by capture and detection of methylated DNA using an ELISA-based method or by separation of methylated and non-methylated nucleotides using liquid chromatography (LC) followed by detection using mass spectrometry or other detection methods (Adamczyk et al., 2023; Tomczyk et al., 2022).

Oxford Nanopore Technologies sequencing enables direct assessment of DNA methylation alongside canonical base sequencing without any additional cost or sample pre-treatment. At higher coverage, it can assess DNA methylation at single-base resolution, and it has recently been shown that global methylation levels can be obtained accurately from very low coverage sequencing, enabling a cost-efficient assessment (Faulk, 2023). Compared with traditional global methylation methods such as ELISA or LC, measuring global methylation on nanopore sequencing offers the advantage of context or region-specific methylation information, a feature especially important in plants where methylation in different cytosine contexts is associated with different biological functions.

Our results indicated that higher coverage (0.1x to 1x) is needed to achieve a comparable precision and accuracy when assessing global methylation in plants using the skimseq approach, compared with vertebrates. The genome sizes of the plant species investigated in the current study are relatively small compared with vertebrates, enabling a cost-efficient assessment of individuals despite the increased coverage required. For example, for *Vitis vinifera*, it would be possible to determine global methylation from 96 multiplex samples sequenced on a single PromethION flow cell.

However, genome size and ploidy vary greatly among plant species (Pellicer & Leitch, 2020; K. Zhang et al., 2019). In addition to having a relatively small genome size, each of the three plant species included in this study is diploid. Further validation of this approach in plant species with larger genomes and higher ploidy levels will be useful to ensure the general applicability of this approach across plant species.

Our data showed that read length affects the accuracy of skimseq methylation assessment. When the *Vitis* data were grouped into sets of differing read length, lower error rates were observed for data with shorter reads. This contrasts with report by (Faulk, 2023) that observed no effect of read length on error rate. Notably, however, the average read length in their dataset was considerably lower (<6kb) while our data also included much longer reads (<10kb up to >50kb). The greater error is likely because very long reads will result in less randomisation during downsampling due to non-independence of datapoints within single reads, especially in regions with highly variable DNA methylation. Indeed, higher accuracy was observed when including only reads shorter than 10kb, particularly in the CHG and CHH contexts, for which error rates dropped from ∼9% to ∼3% and ∼2% at 0.1x coverage, respectively. It is therefore advisable to maintain read length at around or below 10kb, for example, by shearing the DNA before sequencing.

Another factor contributing to error rate is the quality of the reference genome used and the genetic similarity between the reference sample and the test samples. Remapping grapevine sequence data to an in-house reference genome of the same genotype (species, variety and clone) resulted in reduced error rates across all contexts at 1x and 0.1x. We hypothesise that two factors may underlie these observations. Firstly, a more accurate reference assembly will result in fewer read-mapping errors. Secondly, a close genetic match between reference and sample will also reduce misclassification of methylation calls resulting from sequence variation affecting sequence motifs (CG, CHG or CHH) in the reference. As the plant- and variety-specific genome assemblies become more abundant, the accuracy of this approach for plant study can be improved.

Another possible factor contributing to the error rate difference is the heterogeneity or randomness of methylation pattern throughout the genome (which has been termed ‘methylation entropy’) (Xie et al., 2011). Kiwifruit has lower methylation entropy than grapevine, however human and Arabidopsis have similar methylation entropy values which are higher than the two other plant species, but the error rates were lower in human dataset. Therefore, methylation entropy does not appear to explain the error rate differences.

Methylation levels in plants are highly dynamic and can be affected by various environmental and biological factors (Zhang et al., 2018a). Differences in plant global methylation have been observed among tissues and developmental stages (Gao et al., 2019; Shangguan et al., 2020; Teyssier et al., 2008) or after exposure to abiotic stimuli such as heat stress (F. Liu et al., 2023; Yadav et al., 2022), osmotic stress (Antro et al., 2023; Wang et al., 2011), and drought (Antro et al., 2023).

There is currently no published data describing DNA methylation variability between stages of grapevine leaf development or resulting from different sample preservation or extraction methods. Young leaves are the preferred material for grapevine genomic studies due to their amenability to DNA purification techniques. However, no published data could be found comparing methylation levels across different stages of leaf development. Often, plant tissue preservation methods involving the use of liquid nitrogen in the field are impractical. Alternative preservation methods, such as silica gel or dry ice freezing, could offer a practical approach for plant epigenetic studies if proven to preserve DNA methylation information. Our findings showed no measurement effect due to preservation method, both with regards to global methylation as well as across specific genetic features. Similarly, no difference was observed across the two most common approaches for plant DNA purification: SDS and CTAB-based.

Using the skimseq approach, we were able to identify different methylation level of class II transposable elements between two *V. vinifera* varieties. Lastly, the epigenome of grapevine tendril tissue could be distinguished from leaf samples due to elevated CHH methylation, while no significant differences were found between leaf samples. This suggests that a degree of flexibility is possible when collecting young *V. vinifera* leaves for epigenomic studies.

In conclusion, applying the skimseq approach to nanopore sequencing, combined with sample multiplexing appears to be a suitable and cost-efficient method for studying global DNA methylation in plants. Our results show that very long reads are less favoured for measurement precision and accuracy, and therefore DNA shearing, which is known to benefit yield, would also improve the accuracy of methylation measurements for low-coverage sequencing. The quality of the reference genome to which the reads are mapped influences estimate precision and accuracy. Nevertheless, our findings suggest that this method should be broadly suitable as screening tool to study changes in plant global methylation status across developmental stages or due to external stimuli at coverage levels of 0.1x or higher.

## Data Availability

All bioinformatic scripts are available in Github at: https://github.com/yusmiatiliau/Plant_skimseq_methylation. Raw sequence datasets for the grapevine samples have been deposited in ENA under accession number PRJEB78871.

## Acknowledgements

The authors wish to acknowledge the use of New Zealand eScience Infrastructure (NeSI) high performance computing facilities, consulting support and/or training services as part of this research. New Zealand’s national facilities are provided by NeSI and funded jointly by NeSI’s collaborator institutions and through the Ministry of Business, Innovation & Employment’s Research Infrastructure programme. URL https://www.nesi.org.nz. The author(s) also would like to thank Christopher Faulk for the discussion and his insight on the analysis of our data, and TIKI wine (NZ) for contributing to the grapevine tissue samples used in this study. This work was supported by funding from New Zealand Ministry of Business, Innovation, and Employment and from New Zealand Winegrowers.

## Authors contribution

YL, AW and DL conceived the study and co-wrote the manuscript. YL performed all bioinformatics analyses, with AW, SB and SJT contributing reference genome assemblies. YL extracted DNA and performed nanopore sequencing with additional contributions from BV, MPJ, EH, AH. All authors read, edited, and approved the final manuscript.

