## Supplementary Figure 1-5 for "Low-pass nanopore sequencing for measurement of global methylation levels in plants"

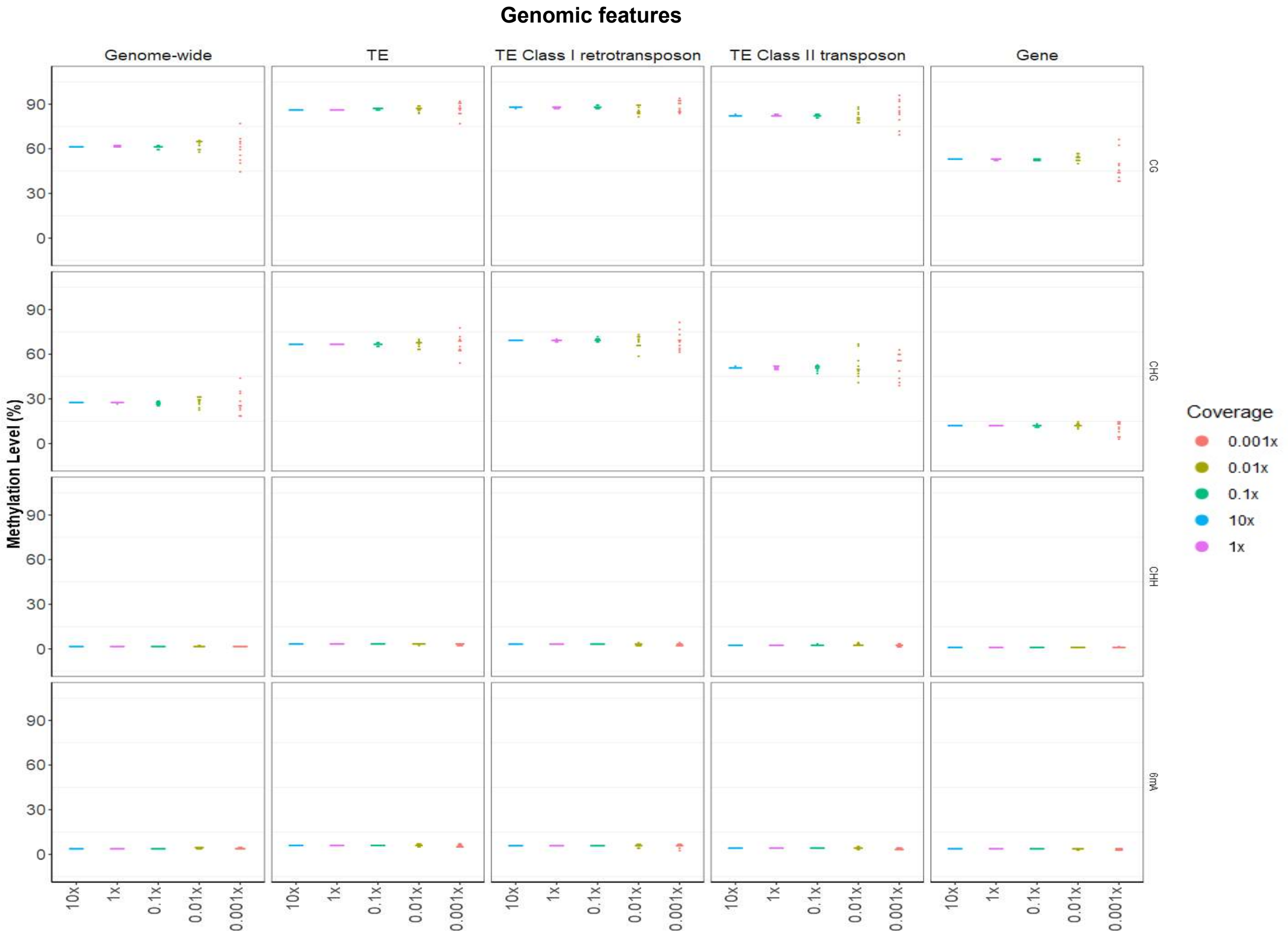


Supplementary Figure 1 Methylation levels of *Vitis vinifera* across different genomic features

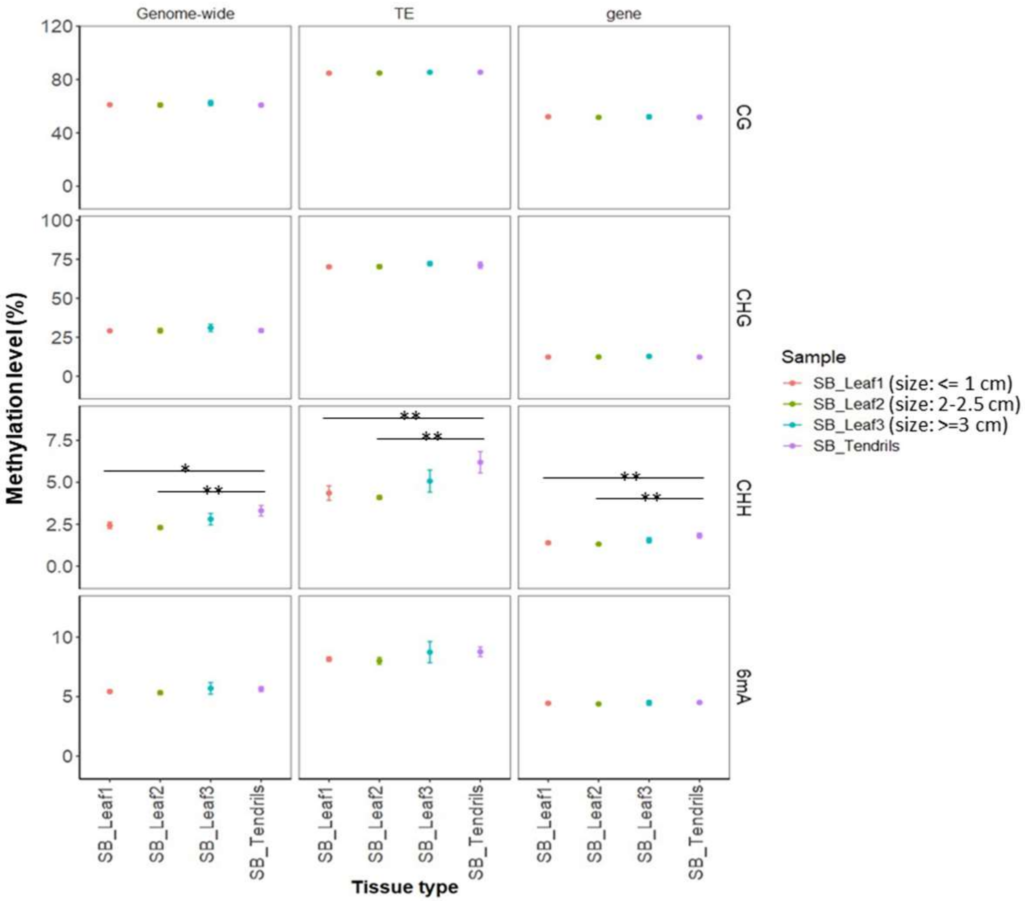


Supplementary Figure 2. Methylation levels (mean ± sd) of different *Vitis vinifera* tissues

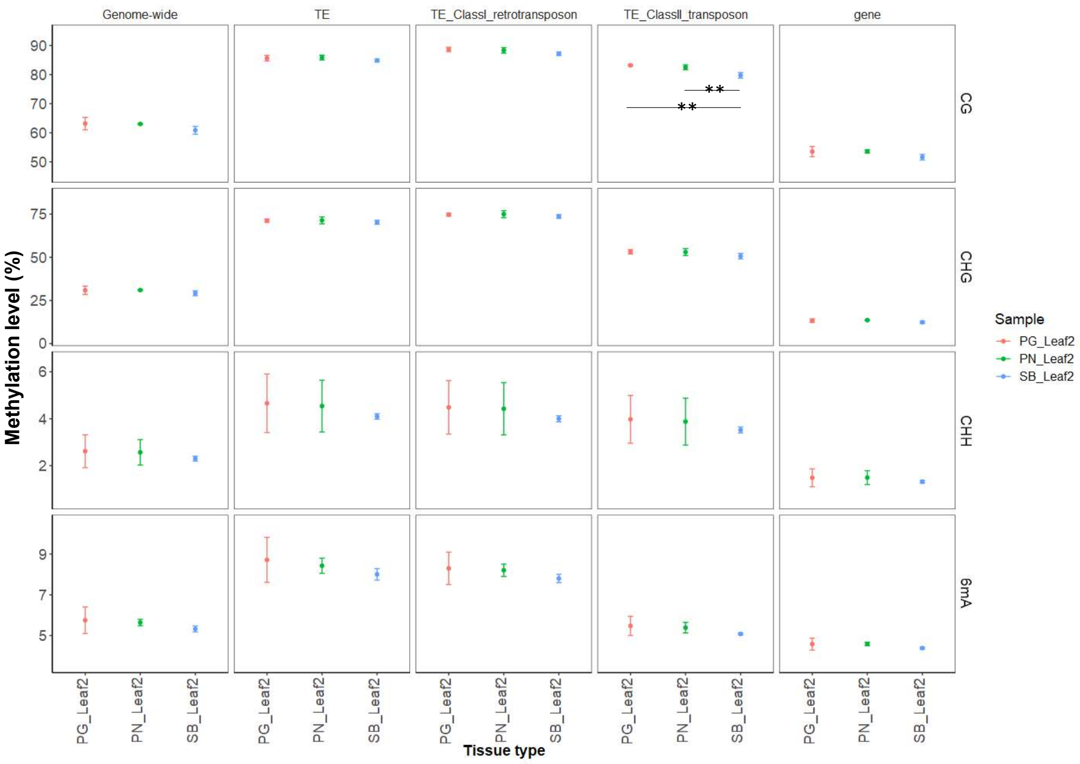


Supplementary Figure 3 Methylation levels (mean ± sd) of different *Vitis vinifera* varieties, Pinot Gris (PG), Pinot Noir (PN), and Sauvignon Blanc (SB)

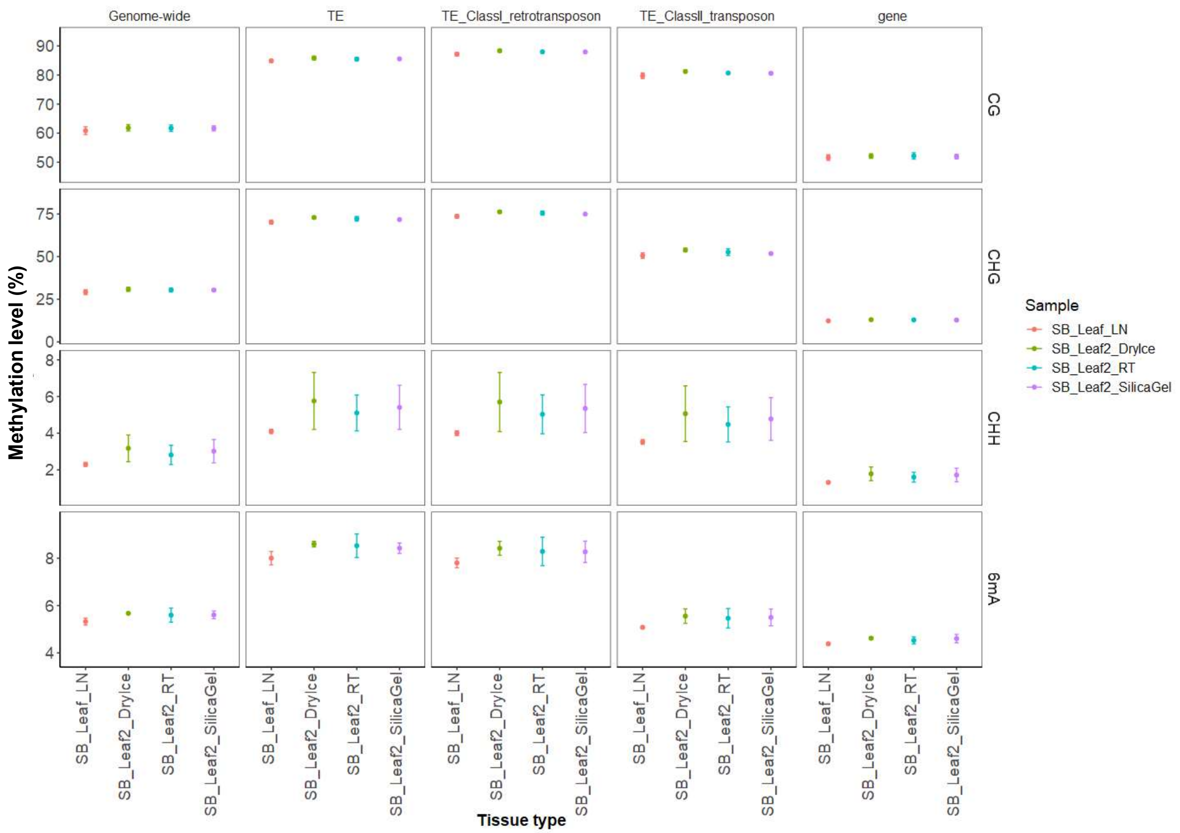


Supplementary Figure 4 Methylation levels (mean ± SD) of *Vitis vinifera* using different sample preservation methods. LN = Liquid Nitrogen, RT = Room Temperature

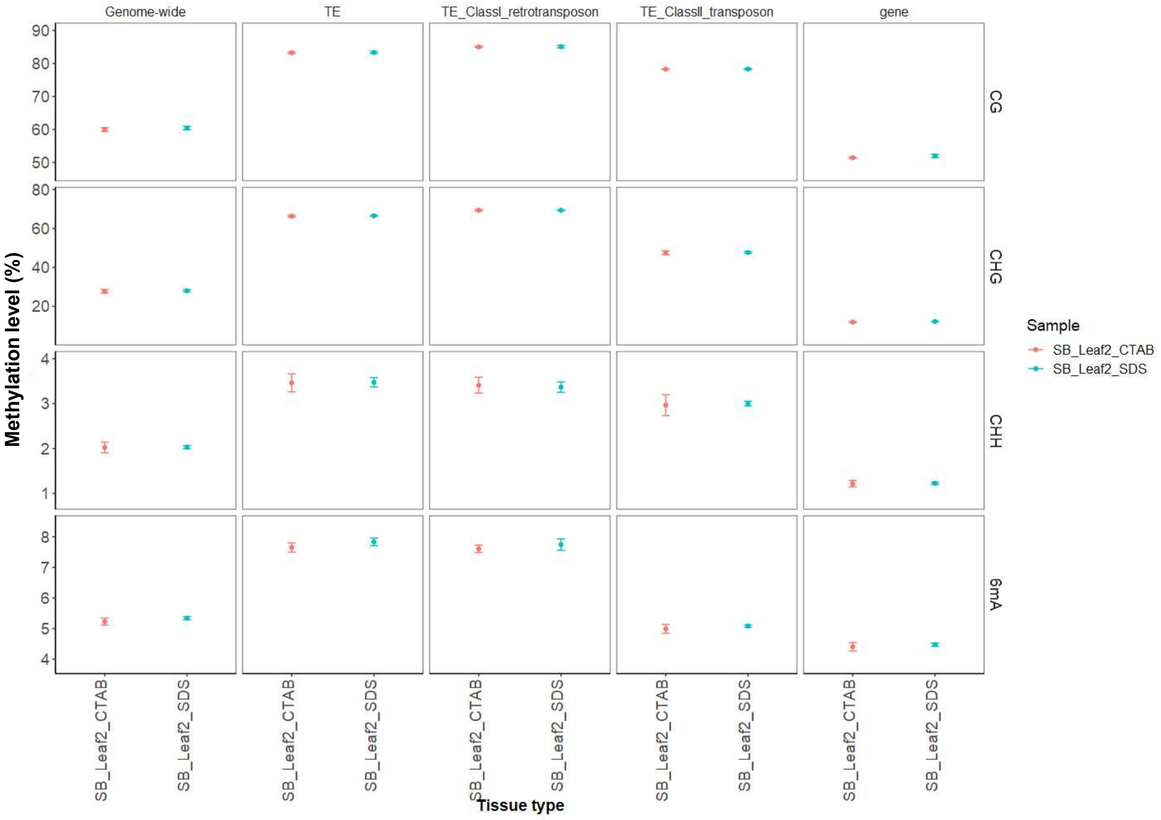


Supplementary Figure 5 Methylation levels (mean ± sd) of *Vitis vinifera* from different DNA extraction methods. CTAB = Cetyltrimethylammonium bromide, SDS = Sodium dodecyl sulfate.
